# Destructive harvest validation of high-throughput measurements show that water use efficiency is unaffected by moderate drought in tobacco

**DOI:** 10.64898/2026.08.28.747842

**Authors:** Samantha S. Stutz, Ron Edquilang, Carl J. Bernacchi, Donald R. Ort

## Abstract

Water-use efficiency (WUE), the ratio of accumulated plant biomass to water lost through transpiration has conventionally been determined using a destructive single-point measurement. Recent advances in high-throughput phenotyping now enable repeated, non-destructive estimation of biomass and WUE. However, these digital measurements must be statistically validated against conventional destructive methods to validate their use as reliable proxies. Therefore, we compared digital biomass determined point clouds produced from multispectral camera scanners with destructive harvests across eight harvests using Samsun tobacco grown under both drought and high-water conditions. WUE efficiency, calculated using the digital biomass estimated from a point cloud and gravimetric water use determinations, were compared to destructive harvest determinations. The coefficient of variation (CV) showed there were no significant differences in digital and destructive measurements for either biomass or WUE. Indicating that digital measurements can be used in place of destructive measurements. Drought plants used significantly less water and were significantly smaller than high-water plants from Harvests 4 through 8. However, there were no significant differences in the ratio of evapotranspiration to leaf area or WUE, indicating that drought plants were simply smaller and used less water than the high-water plants. This work validates that estimating plant biomass from a digital point coupled with continuous gravimetric determination of water use provides a reliable nondestructive measure of WUE in high-throughput measurements across the full plant life cycle.

**PLAIN LANGUAGE SUMMARY:** We grew tobacco plants under either a drought or high-water treatment and harvested a portion of the plants every few days for a total of eight harvests. Throughout the experiment, we collected 3D images of the plants and continuously measured pot weight to track plant growth and water use across different developmental stages. Destructive biomass served as the gold-standard measurement. We then compared biomass and water-use estimates generated from the digital measurements with the destructive measurements. The digital approach provided accurate estimates of plant biomass and water use while requiring little hands-on labor and no plant destruction. These nondestructive methods could help plant breeders identify water-efficient plants earlier in the breeding process, accelerating the development of crops that use water more efficiently.

## 1 INTRODUCTION

Photosynthesis requires open stomata to permit CO_2_ diffusion into the leaf that in turn leads to water loss. Limiting water loss while maximizing plant growth has been a target for improving resilience and productivity for both C_3_ and C_4_ crops for over a century (Boyer 1982), particularly given that agriculture accounts for 80% of global fresh water use (FAO). Water-use efficiency (WUE) is the ratio of carbon gain to water loss commonly approximated using intrinsic WUE, which relates the ratio of net CO_2_ assimilation to stomatal conductance, whereas, instantaneous WUE is the ratio of net photosynthesis to transpiration rate. Both intrinsic and instantaneous WUE can be quickly and easily estimated using gas exchange systems. However, in grapes (*Vitis vinifera*), instantaneous WUE varied with time of day, season, and canopy position (Medrano et al., 2015), so conclusions based on this metric are context dependent. In contrast, whole plant WUE, the ratio of total biomass accumulated to total water consumption (Briggs and Shantz 1913), is more difficult to quantify because measuring lifetime water use is a labor-intensive, destructive and single-time point measurement. However, the advent of high-throughput phenotyping technologies potentially enable non-destructive, repeated estimates of plant biomass and water use through time. Digital WUE estimated from high-throughput phenotyping enable non-destructive estimates of plant biomass and water use at multiple times throughout the growing season and across the life of an individual plant. Estimating WUE of an individual across its life cycle is a valuable measurement for breeding crops more resilient to climate change. Water availability is particularly critical during specific developmental stages, such as the reproductive phase, when processes like pollination and grain set are highly sensitive to drought stress. For example, crops often exhibit greater yield reductions under water deficit during flowering than during vegetative growth (Blum, 2009; Farooq et al., 2009, Farooq et al., 2019). However, quantifying how water use efficiency (WUE) varies across developmental stages remains challenging, largely because whole-plant WUE is difficult to estimate repeatedly over time with destructive biomass measurements.

High-throughput plant phenotyping platforms enable repeated nondestructive estimation of traits such as above ground plant biomass, and leaf area. When these systems also measure changes in plant and pot mass, they can quantify water use through time. Together, repeated estimates of biomass and water use create the opportunity to quantify whole-plant WUE throughout development rather than only at final harvest. However, the utility of these digital measurements depends on their validation against conventional destructive (reference) measurements. McGrath et al. (2024) and Siebers et al. (2024) recently developed a framework for evaluating agreement between digital and destructive measurements by considering bias and variance rather than relying on correlation alone. When digital and destructive measurements differ in units, dimensionless statistics such as coefficient of variation (CV) can be used to compare their relative variability (Feltz and Miller, 1996; Marwick and Krishnamoorthy, 2019). To complement this, regression approaches can be used to calibrate digital against destructive measurements and test their ability to predict conventional estimates in data independent of model development (Linnet, 1990; Meacham-Hensold et al., 2019).

With proper calibration against destructive measurements, phenotyping platforms that weigh and image plants allow repeated study of plant growth and water use at the organismal level. Phenotyping platforms that incorporate gravimetric measurements can measure total evapotranspiration on a daily or shorter timescales depending upon the measurement frequency. This temporal resolution provides an opportunity to determine how plant growth, water use, and ultimately WUE respond dynamically to water limitation. Water-stress affects species differently depending upon when the drought-stress occurs, and the duration of the stress; however, drought-stressed plants typically have less biomass compared to non-water-stressed plants (Farooq et al., 2009; Blum, 2011; Khamis et al., 2025), which is usually accompanied by a decrease in transpiration (Farooq et al., 2009). In tobacco, short-term water deficits reduce plant biomass and leaf area with water use declining with water limitation (Ryan et al., 2013; Turc et al., 2024). Instantaneous WUE often increases under water deficit conditions because transpiration and stomatal conductance decline proportionally more than photosynthesis (Hatfield & Dold, 2019). However, this response does not necessarily translate into greater whole plant WUE, which can increase, remain the same or decrease under drought (Tomas et al., 2012). This discrepancy may be the result of high rates of night transpiration, high cuticular conductance (Escalon et al., 2013), and/or other water losses that are not captured by instantaneous leaf-level measurements. Repeated measurements of biomass and water use therefore provide a means to determine whether drought-induced reductions in plant size and transpiration ultimately alter whole-plant WUE over the course of development.

Therefore, the objectives of this study were to (1) compare biases and variance between digital and destructive measurements of biomass and WUE to determine whether nondestructive high-throughput phenotyping can reliably estimate whole-plant WUE and (2) determine how biomass, water use, and WUE change in tobacco subjected to moderate drought stress. Tobacco was selected as a model phenotyping species because of its relatively simple architecture, and short life-cycle. We hypothesized that the digital estimates of biomass and WUE would exhibit minimal bias and variability comparable to destructive, supporting their use as non-destructive proxies. We further hypothesized that moderate drought would reduce biomass and water use while increasing whole-plant WUE.

## 2 MATERIALS AND METHODS

### 2.1 High Throughput Phenotyping Facility (HTPF)

The University of Illinois High Throughput Phenotyping Facility (HTPF) in Champaign, IL, USA consists of a Phenospex FieldScan® System (Phenospex, Heerlen, Netherlands) with 384 FieldScales (Phenospex) arranged into 32 blocks with 12 FieldScales in each block (Fig. S1). Each FieldScale was irrigated using the DroughtSpotter (Phenospex) automated watering system. The FieldScales were scanned with two pairs of PlantEye F500 multispectral 3D scanners (Phenospex). A complete scan of all 384 FieldScales required approximately 75 min. The validation experiment used wild-type *Nicotiana tabacum* (cultivar Samsun). The experiment was conducted from October through November 2023 (Table 1).

**Table 1.** Important dates for sowing, transplanting, and harvesting for the tobacco experiment. Days after transplanting start at zero the day all the seedling were transplanted into the pots and placed on the FieldScales.

| Days after sowing | Days after transplanting | Event | Number of plants harvested | Date (yyyymmdd) | Plant development | Notes |
| --- | --- | --- | --- | --- | --- | --- |
| 0 | - | Sowing | - | 2023-09-26 |  |  |
| 14 | - | Transplanting to float trays | - | 2023-10-10 |  |  |
| 27 | 0 | Transplanting into pots, start experiment | 0 | 2023-10-23 |  |  |
| 36 | 9 | Harvest 1 | 48 | 2023-11-01 |  |  |
| 38 | 11 | Harvest 2 | 48 | 2023-11-03 |  |  |
| 42 | 15 | Harvest 3 | 48 | 2023-11-07 |  | Start of drought experiment |
| 45 | 18 | Harvest 4 | 48 | 2023-11-10 |  |  |
| 49 | 22 | Harvest 5 | 47 | 2023-11-14 |  | One pot never received Osmocote. Small yellow, HW plant excluded from data. |
| 52 | 25 | Harvest 6 | 48 | 2023-11-17 | 4 H plants buds<br>3 D plants buds |  |
| 56 | 29 | Harvest 7 | 47 | 2023-11-21 | 22 H plants buds<br>16 D plants buds | One FieldScale not properly calibrated. HW plant excluded from data. |
| 63 | 36 | Harvest 8 | 48 | 2023-11-28 | 2 H plants buds<br>22 H plants flowers<br>3 D plants buds<br>21 D plants flowers |  |

### 2.2 Plant propagation

Tobacco seeds were sown in a single azalea pot with a diameter of 15 cm on 26 September 2023. The pot was placed in a growth chamber (Conviron, Winnipeg, Manitoba, Canada) at 700 μmol quanta m^-2^ s^-1^ and 65% relative humidity under a 15 h/9 h day/night photoperiod and 25°C/23°C day/night temperatures at the HTPF. On 10 October 2023 (14 DAS) seedlings were transplanted to float trays and moved to the greenhouse. The seedlings were transplanted on 23 October 2023 (27 DAS) into the pots on the FieldScales.

Mean HTPF conditions were 28.0°C/27.0°C day/night, and 58.0%/55.3% day/night relative humidity. The greenhouse shade cloth was deployed during the entire experiment to reduce the light gradient that naturally occurs in the greenhouse. One hundred forty LED light sources (ELEXIA model LX6XX; Heliospectra; Gothenburg, Sweden) provided supplemental lighting for the experiment. The light sources provided an average irradiance of 500 μmol quanta m^-2^ s^-1^. Including natural lighting, the light, average irradiance during the experiment was approximately 1000 μmol quanta m^-2^ s^-1^.

### 2.3 Experimental layout

We used a randomized complete block design (Fig. S2). The 32 blocks were assigned among eight harvests, with four blocks per harvest (Fig. S2). With the exception of Harvests 5 and 7, each harvest included 48 plants: 24 high and 24 drought-treated plants (Table 1). A high-water plant in Harvest 5 that did not receive fertilizer and a high-water plant in Harvest 7 with a malfunctioning FieldScale were excluded from the data (Fig. S2, Table 1). Each block had six high-water and six drought plants.

### 2.4 Field capacity (FC) estimates and transplanting

Field capacity (FC), the amount of water held by a soil after excess water has drained, was estimated for 6.5 L Classic 600 pots (Nursery Supply Co. Louisville, KY, USA) with BM6 potting soil (Berger, Saint-Modeste, Quebec, Canada). Soil was prewetted in a large bin and placed in the Classic 600 pots. Pots were placed on the FieldScales and watered to drainage (field capacity). The maximum pot mass immediately before drainage-associated mass loss was taken as field capacity. Mean FC across pots was 4175 g. Variation among pots likely reflected differences in soil compaction among operators and initial water content among batches of prewetted soil. Irrigation was controlled in HortControl (Phenospex) using deviation mode. In deviation mode, the DroughtSpotter applied water when pot mass deviated 5% below the set target mass of 2733.8 g (65% FC) for the high-water treatment or 1708.6 g (41% FC) for the drought treatment.

At 27 days after sowing (DAS) seedlings were transplanted from the float trays into pots with two tablespoons of Osmocote 20-20-20 slow-release fertilizer (ScotsMiracle-Gro, Marysville, OH, USA), then placed on the FieldScales. The pots were allowed to dry to their set FC before they were watered again. We considered Harvest 4, the onset of the drought treatment phase because by that harvest more than 50% of drought-treated pots had reached their irrigation threshold and been rewatered. Harvest 1, 2, and 3 were used to generate the regression for digital biomass. Plants were scanned with the PlantEye every 2 h throughout the experiment. Plants were harvested twice weekly during exponential growth at 9, 11, 15, 18, 22, 25, 29, and 36 days after transplanting (DAT). All plants had buds or were flowering by the last harvest (Table 1).

### 2.5 Soil evaporation

To measure soil evaporation, forty-eight Classic 600 pots were filled with prewetted soil and watered to the average FC as performed above. Thirty pots were divided equally between the high-water and drought treatments (n = 15 per treatment), and the soil was allowed to dry to the respective treatment target before rewatering. Average soil evaporation was calculated for each harvest. The remaining 18 pots were covered with filter paper that was the approximate size of the average tobacco plant when the plants were transplanted into the pots on the FieldScales, ∼13 cm diameter. The filter paper served as a mock plant to estimate soil evaporation during Harvests 1-3, when plants were small and before the canopy covered the soil surface. For Harvests 1-3, mean evaporation from the mock-plant pots in each treatment was subtracted from the cumulative gravimetric water loss of each planted pot in the corresponding treatment. For Harvests 4-8, the cumulative soil evaporation for Harvest 3 (1425 g for drought and 1444.99 g for high-water) were used as the soil-evaporation correction for each treatment. This approach assumed that additional soil evaporation after Harvest 3 was small because, on average, plant canopies covered the soil surface during Harvests 4-8 and a surface crust formed as the soil dried. These evaporation corrections were applied to the gravimetrically determined water loss term used in the WUE calculations.

### 2.6 Harvest

Plants were harvested following a PlantEye scan; the shoot was cut at the soil level. Smaller plants were weighed intact; for larger plants, leaves were removed and stems were sectioned before weighing. The number of leaves, axillary branches, and the presence of flowers or buds were noted. Leaf area was measured using a LI-3000 leaf area scanner (LI-COR Biosciences, Lincoln, NE, USA). After the leaf-area measurement, the plant was weighed again. Following the harvest, the plants were placed in a drying oven at 60°C for two weeks and weighed to obtain dry biomass.

### 2.7 Digital biomass analysis

PlantEye scans were reviewed in Phenospex, HortControl for quality control. Individual plants were inspected to confirm that the plant was contained within its scan area, and that adjacent leaves or soil were not included in the scan. No scans required manual exclusion or cleanup based on these criteria.

### 2.8 Estimates of whole plant water-use efficiency

Whole plant water-use efficiency, was calculated from transplanting to each harvest using the same cumulative gravimetric water use denominator for both reference and digital estimates as:

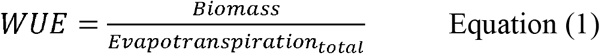

For destructive WUE, the aboveground wet mass was used to calculate WUE to be consistent with digital measurements which were taken on turgid plants. Digital WUE used the digital biomass estimate from the final PlantEye scan before harvest. The cumulative water-use denominator was identical for both WUE estimates and was calculated gravimetrically according to Peters (2023) as:

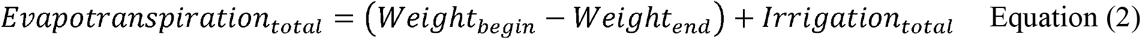

where, the Weight_begin_ was the initial mass of the pot, soil, dripper and plant and Weight_end_ was the corresponding mass at harvest. Irrigation_total_ is the total amount of water added by the DroughtSpotter and recorded by the Phenospex over the course of the experiment. The soil-evaportation correction described in Section 2.5 was then applied to the gravimetric water loss estimate used for WUE.

### 2.9 Coefficient of variation (CV) estimates

Because destructive and digital estimates of biomass and WUE use different units, their absolute values were not directly comparable. Relative variability between methods was therefore compared using the coefficient of variation (CV), following the procedure in Siebers et al. (2024):

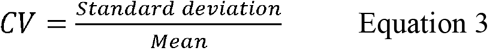

For each treatment-by-harvest combination, the mean and standard deviation were calculated from 24 plants except for high-water plants in Harvests 5 and 7, which included 23 plants. CVs were calculated separately for the drought and high-water treatments at each harvest. Equality of CVs between digital and reference measurements of biomass and WUE were tested according to Feltz and Miller (1996) using the asymptotic test implemented the cveqality R package (Marwick and Krishnamoorthy, 2019).

### 2.10 Predictive models for destructive biomass and destructive WUE

Blocks within each harvest were randomly divided between a training dataset (50%) and an independent test dataset (50%) to determine whether digital measurements could accurately predict reference measurements. Deming regression, an errors-in-variables approach that accounts for measurement error in both variables (Linnet, 1990), was used to derive calibration relationships between digital and reference measurements. Calibration equations derived from the training data were applied to digital measurements in the test dataset to generate predicted reference values. Predicted reference biomass and WUE were plotted against the corresponding reference measurements in the test dataset. Agreement between predicted and reference measurements was evaluated relative to the 1:1 line and by linear regression.

### 2.11 Statistical analysis

We used R (version 4.4.2, R Development Core Team 2024) for all statistical analyses. We used one-way ANOVAs to compare between treatments and among harvests for total evapotranspiration, reference and digital WUE, destructive and digital biomass, and the ratio of evapotranspiration to destructive and digital leaf area.

## 3 Results

### 3.1 Association between digital and reference measurements of biomass and water-use efficiency (WUE)

Digital biomass and destructive biomass were strongly positively related across all harvests (R^2^ = 0.95; Fig. 1). Plant size also increased across successive harvests, with smaller plants in earlier harvests and larger plants in later harvests. In contrast, digital WUE and reference showed little relationship (R^2^= 0.05; Fig. 2).

**Figure 1.**
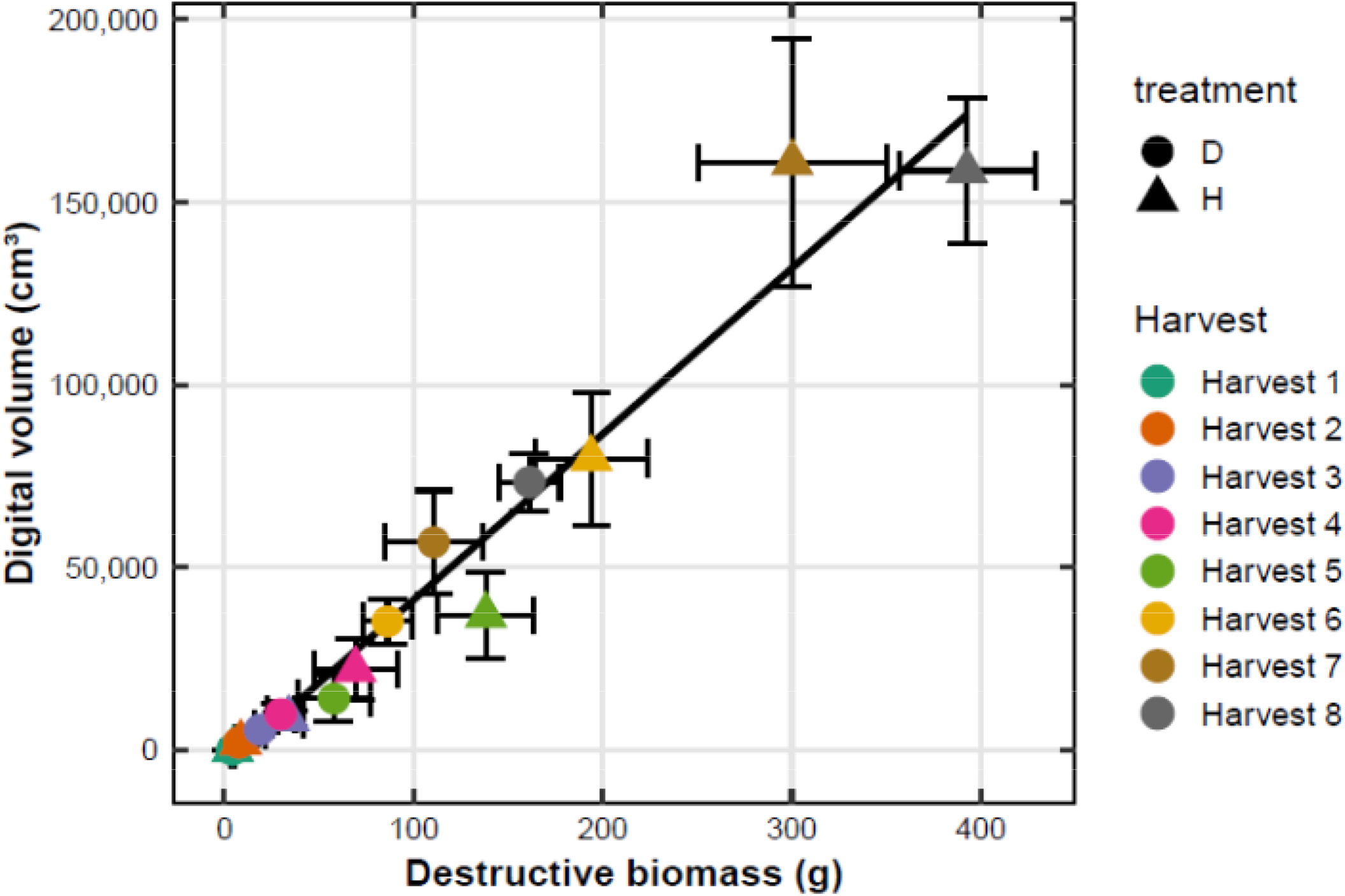
Correlation between digital biomass and destructive biomass. Digital biomass estimated from the PlantEye is correlated with destructive biomass. Tobacco plants were grown under drought conditions (D—41% soil field capacity) and high-water (H—65% soil field capacity). Each point represents the mean of four blocks ±1 SD.

**Figure 2.**
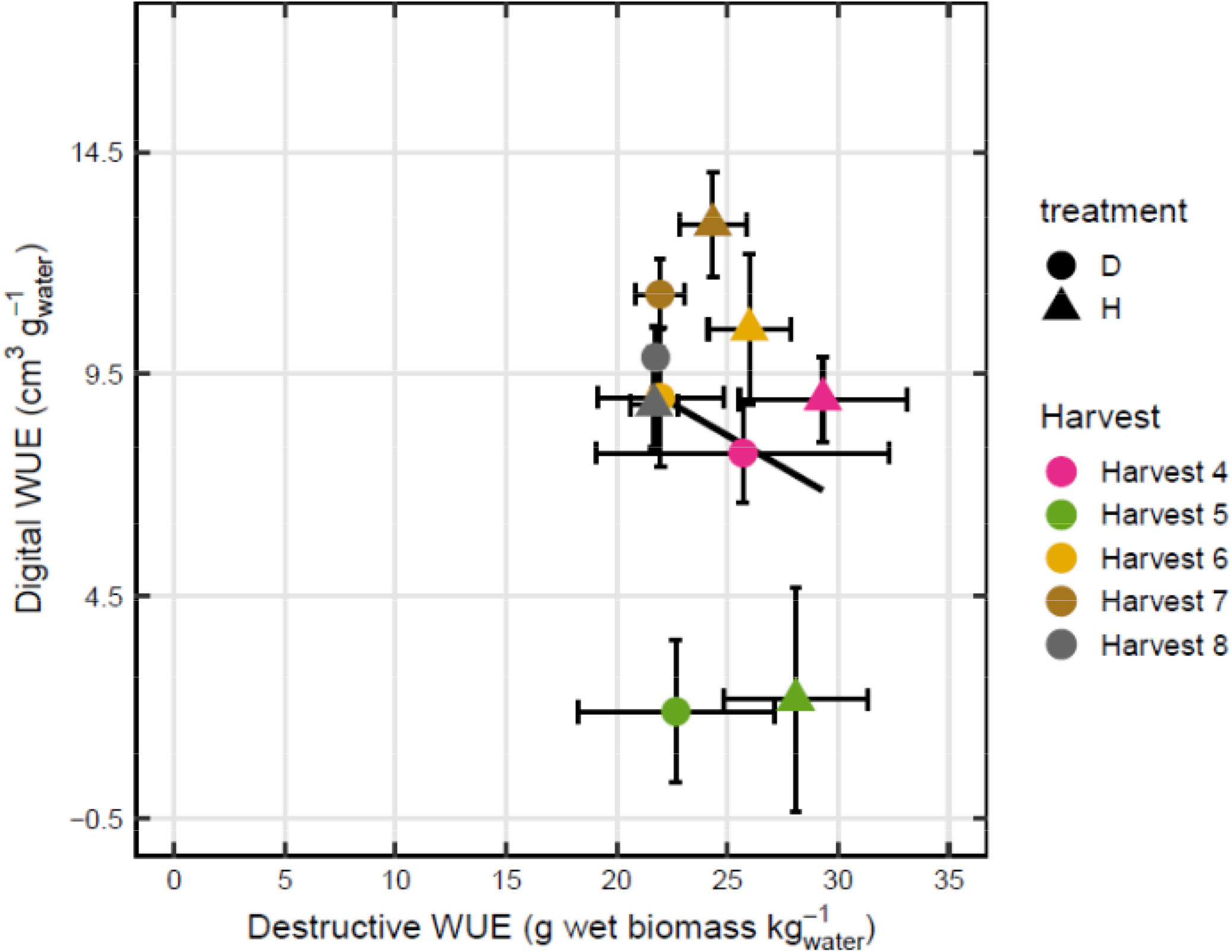
Digital WUE plotted against destructive WUE for Harvests 4-8, the duration of the drought experiment. Tobacco plants were grown under drought conditions (D—41% soil field capacity) and high-water (H—65% soil field capacity). Each point represents the mean of four blocks ±1 SD.

### 3.2 Coefficient of variation

Because digital and reference biomass and WUE were expressed in different units, the coefficient of variation (CV) was used to compare relative variability between methods. The CVs did not differ significantly between digital and destructive biomass for either the drought (*p* = 0.440) or high-water treatment (*p* = 0.81; Table 2). Likewise, the CVs did not differ significantly between digital and reference WUE for either the drought (*p* = 0.94) or high-water treatment (*p* = 0.19; Table 2).

**Table 2.**
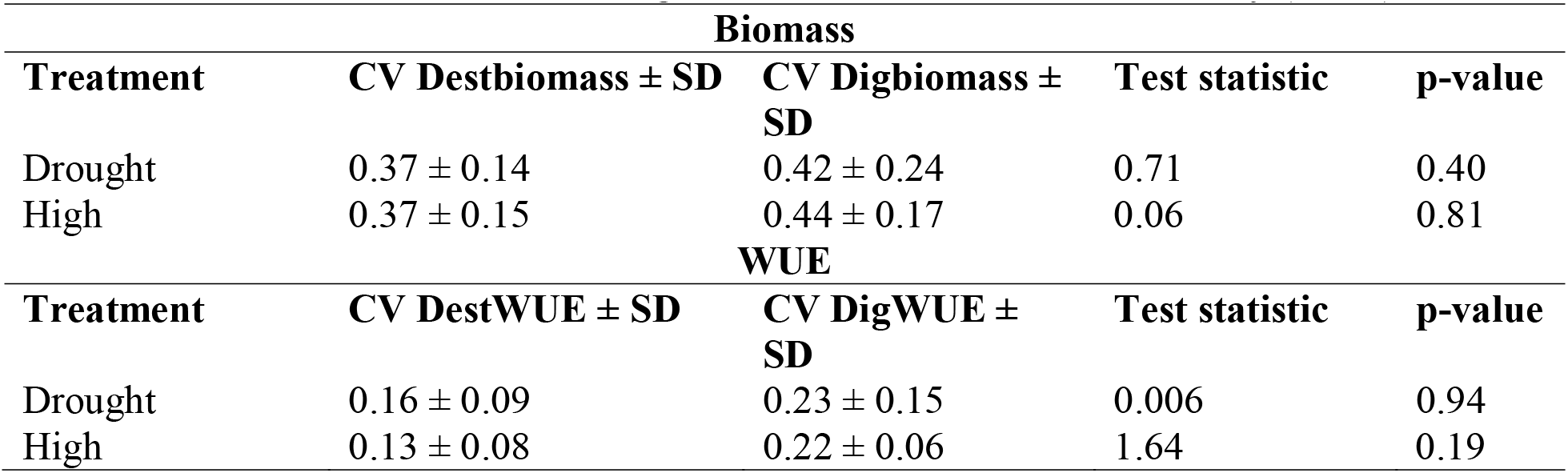
Coefficient of variation for digital biomass and water-use efficiency (WUE).

### 3.3 Prediction of reference measurements from digital measurements

Data were divided into a training and a test dataset, and a Deming regression fit to the training dataset was used to predict destructive biomass and WUE in the independent test dataset. The resulting predictions were expressed in the same units as the corresponding reference measurements, allowing direct comparisons between methods. Predicted destructive biomass closely followed the 1:1 line for both treatments through Harvest 6, whereas deviations increased for the larger high-water plants in Harvests 7 and 8 (Fig. 3A). The bias plot similarly showed that most smaller plants fell within the limits-of-agreement, while several of the largest plants fell outside these limits; plants in Harvest 7 tended to be overestimated while plants in Harvest 8 tended to be underestimated with the PlantEye (Fig. 3B). The F-test indicated no significant difference between predicted and destructive biomass (*p* = 0.58).

**Figure 3.**
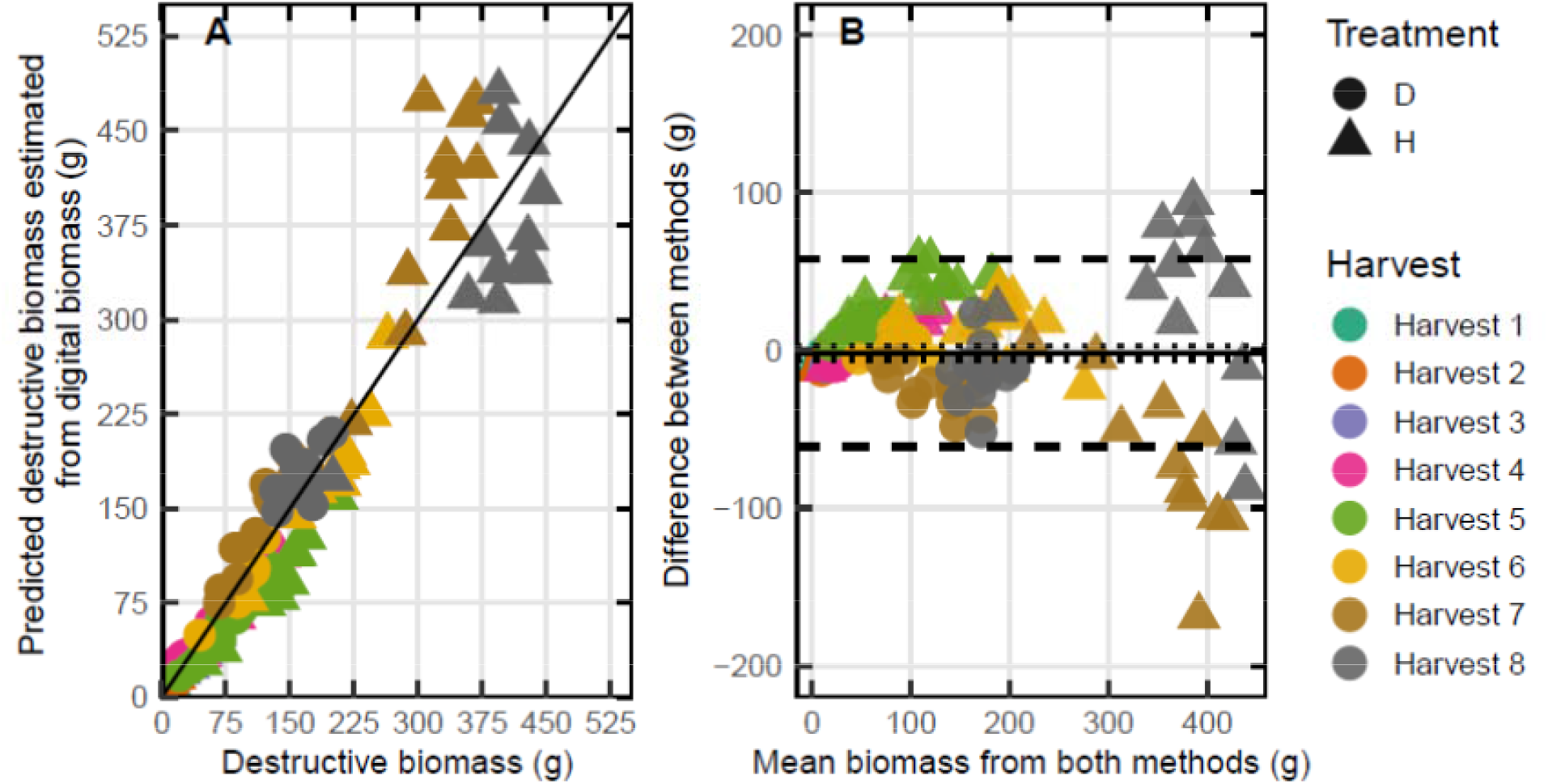
A) Linear regression estimating destructive biomass using the digital volume estimates. Half of the data were used to generate the Deming Regression model. The data in this figure show the test data (half of the data that the Deming Regression was applied to). The 1:1 line indicates a high correlation. **B)** Bias plot with mean bias (solid line), standard error of the mean (dotted line), limits of agreement (dashed line).

WUE varied over a relatively narrow range across treatments and harvests, with many observations near 20 g fresh biomass kg^-1^ (Fig. 4A). Predicted WUE values for Harvests 4 and 5, deviated more from the 1:1 line than those for Harvests 7 and 8 (Fig. 4A). Most observations fell within the limits of agreement in the bias plot (Fig. 4B). The F-test indicated no significant difference between predicted and reference WUE (*p* = 0.92).

**Figure 4.**
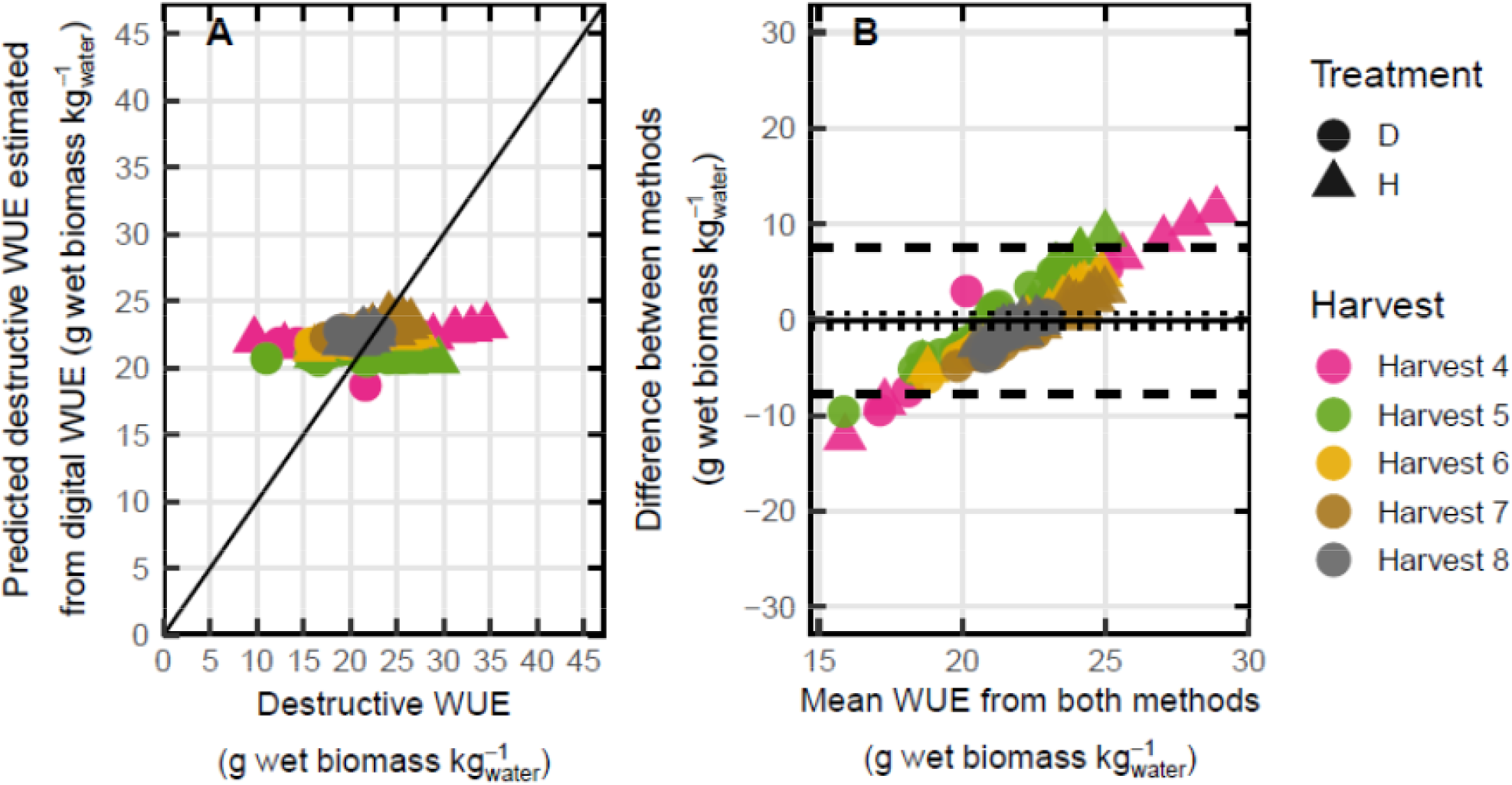
A) Digital WUE estimated using the Deming regression plotted against destructive WUE for the test data set. Estimates of destructive WUE from digital WUE indicate there is a good fit. The data in this figure show the test group of data. The black line is a 1:1 line plotting destructive to digital WUE. These values were obtained by using the Deming Regression model applied directly to the digital WUE. **B)** Bias plot with mean bias (solid line), standard error of the mean, limits of agreement (dashed line).

### 3.4 Tobacco growth under reduced water

Biomass increased across harvests in both treatments. Destructive measurements showed that high-water plants were significantly larger than drought plants in Harvests 5-8 (*p* < 0.001), and digital measurements showed the same pattern over Harvests 5-8 (*p* < 0.01; Fig. S3). At Harvest 8, high-water plants were 144% and 116% larger than drought plants based on destructive and digital measurements, respectively. Digital biomass increased from Harvests 1-6 in both treatments but decreased slightly in high-water plants between Harvests 7 and 8 (Fig. S3).

### 3.5 Evapotranspiration under reduced water

Total evapotranspiration was significantly lower in drought than high-water plants in Harvests 5-8 (*p* < 0.05; Fig 5A). No significant differences in total evapotranspiration were detected between treatments in Harvests 1-4. By Harvest 4, plant leaves covered most of the soil surface reducing the exposed soil area. The average hourly rate of evapotranspiration for high-water and drought plants in Harvest 7 was calculated over a 43 h period (23-25 DAT). Daytime evapotranspiration in high-water plants was more than twice that of drought plants and was occasionally approximately threefold higher (Fig. 5B). Nighttime water loss occurred in both treatments and was greater in high-water plants; nighttime rates were approximately 10% of daytime rates in both treatments.

**Figure 5.**
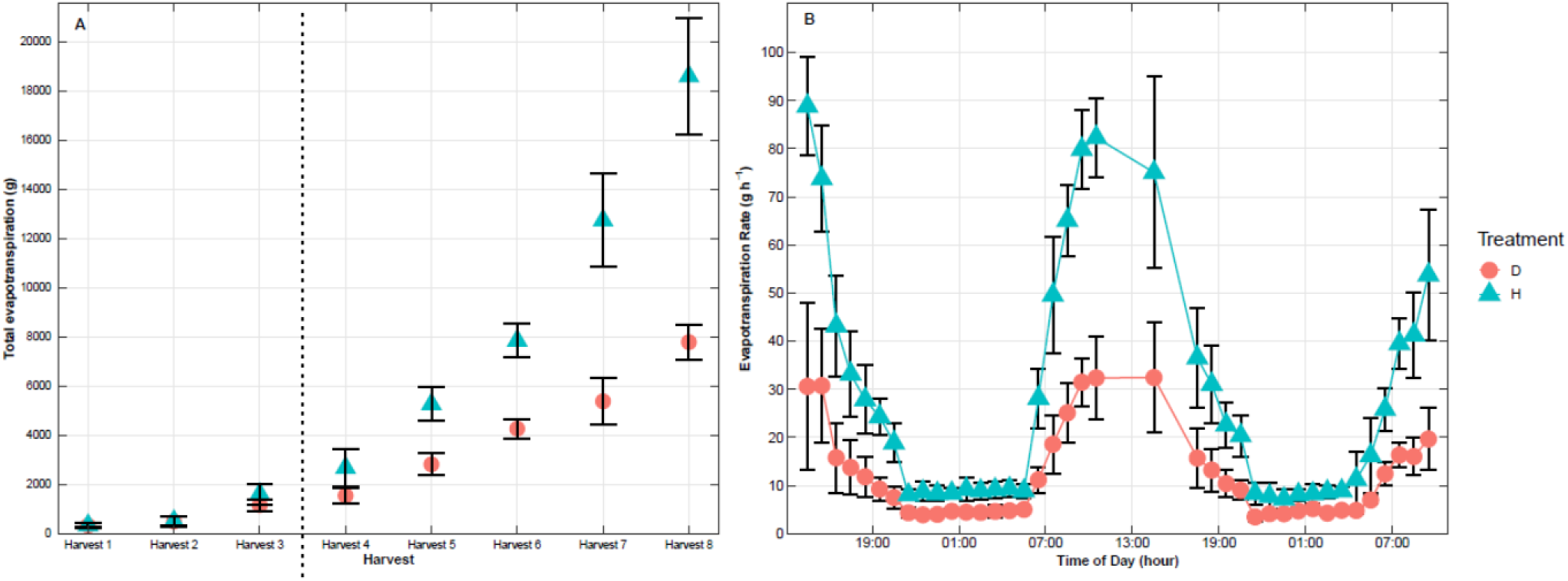
A) Total evaportranspiration from the start of the experiment to the date of harvest. Each point represents mean N=4 ± SD. **B)** Water loss over 43 hours from 20231115 14:30 to 20231117 09:30 (23-25 DAT), for the drought (D—41% soil field capacity) and high-water (H-65% soil field capacity). N is the mean rate of ET for between 1 and 4 blocks error bars ±1 SD.

The ratio of evapotranspiration to destructive leaf area was highest in Harvests 1 and 2 and declined by more than half by Harvest 3. No significant differences between treatments were detected at any harvest (Fig. S4A). In contrast, the ratio of evapotranspiration to digital leaf area increased across harvests and was five- and six-fold greater in Harvest 8 than Harvest 1, for drought and high-water plants, respectively (Fig. S4B). The treatments differed significantly at Harvest 8 (*p* < 0.001).

### 3.6 Whole-plant WUE under reduced water

There were no significant differences between treatments or among harvests for destructive WUE (Fig. 6A). Reference WUE in drought plants increased by ∼14% from Harvest 4 to Harvest 8, whereas destructive WUE in high-water plants decreased by ∼25% over the same interval. By Harvest 8, both treatments had a destructive WUE of ∼22 g wet weight kg water^-1^. Digital WUE decreased in both treatments between Harvests 4 and 5, and increased at Harvest 6; no significant differences in digital WUE were detected between treatments within any harvest (Fig. 6B). Harvest 5 differed significantly from Harvests 6 and 7 for both treatments (*p* < 0.01).

**Figure 6.**
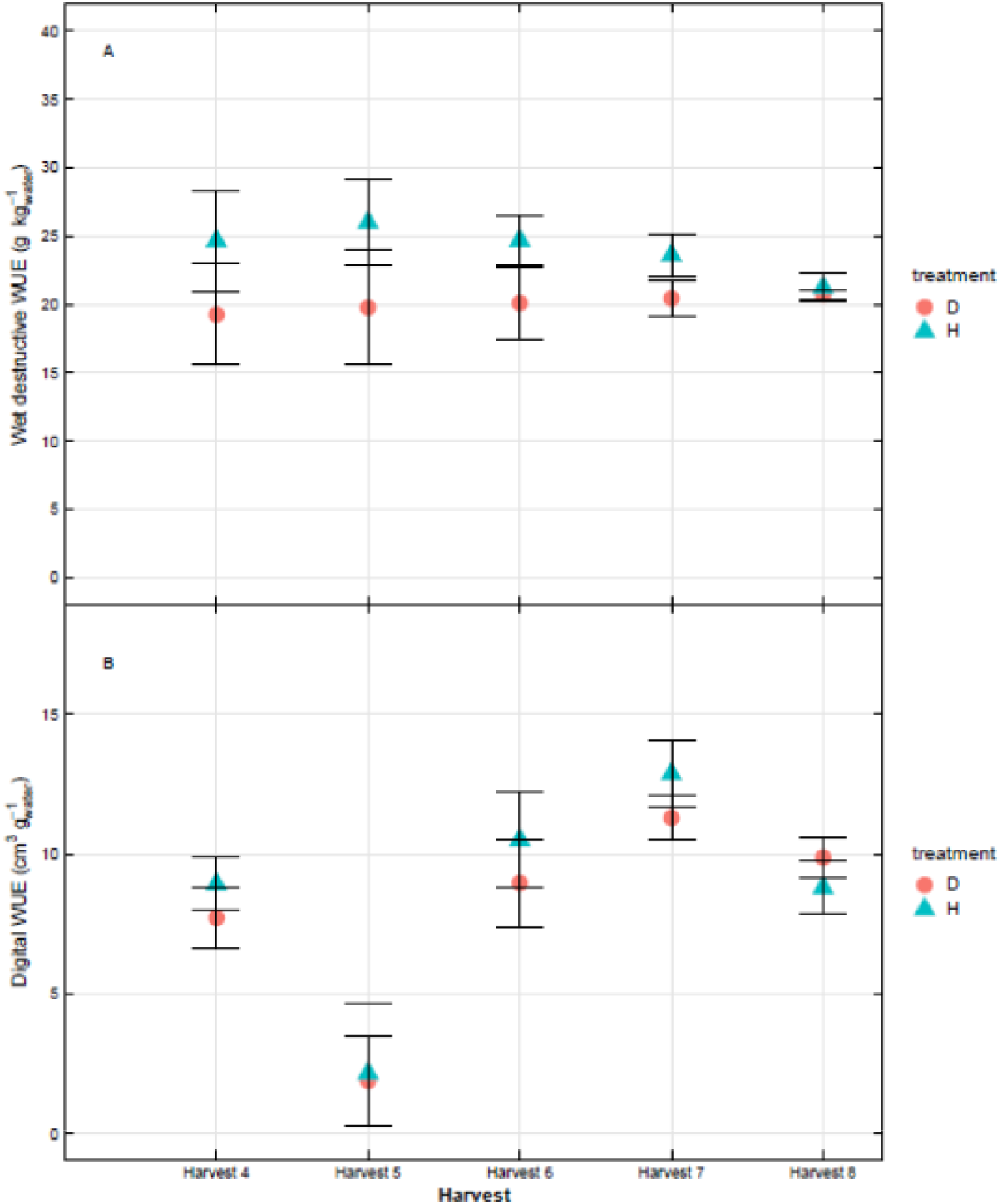
A) Destructive WUE and **B)** Digital WUE for Harvests 4-8, the duration of the drought experiment. Tobacco plants were grown under drought conditions (D—41% soil field capacity) and high-water (H—65% soil field capacity) error bars represent ±1 SD.

## 4 Discussion

High-throughput phenotyping enables rapid, nondestructive, and repeated measurements of individual plants throughout development with minimal manual effort. However, digital measurements must be evaluated against the convention destructive measurements to establish their precision and accuracy. The coefficient of variation (CV) provides a dimensionless measure of relative variability that can be compared between measurements expressed in different units (Marwick and Krishnamoorthy, 2019; McGrath et al., 2024). The CVs did not differ significantly between digital and destructive measurements of biomass or water-use efficiency (WUE), indicating similar relative variability between methods (Table 2). Deming regression further showed that digital measurements predicted destructive biomass well over much of the measured size range, although prediction accuracy declined for the largest plants (Fig. 3). In contrast, prediction of destructive WUE from digital WUE was weaker (Fig. 4), indicating that validation of WUE was less conclusive than validation of biomass. Moderate drought reduced plant biomass and water use but did not significantly alter whole-plant WUE.

### 4.1 Performance and limitations of digital measurements

New technology and high-throughput measurements of plant traits are appealing because of the ease and speed of data collection. However, their performance should be quantified relative to established destructive measurements before they are used as substitutes for conventional methods (McGrath et al., 2024). The coefficient of variation (CV) test made it possible to compare relative variability between digital and destructive measurements of biomass and WUE despite differences in measurement units. The absence of significant differences in CV indicates that the methods had similar relative variability, but does not by itself demonstrate equivalence or absence of bias. Destructive biomass can be measured only once for an individual plant, and variability in destructive measurements can be minimized by using consistent harvest procedures. Digital measurements also contain sources of variability; the larger CVs observed for some digital measurements could reflect leaf movement during scanning, overlapping leaves in larger canopies, or other sources of measurement error. Averaging multiple successive scans may reduce random variation in digital measurements, although this should be tested directly. For digital measurements to be directly interpretable, calibration to the units of the destructive measurement is useful. Splitting the data into training and test sets allowed us to generate a Deming regression relating digital biomass to destructive biomass and then evaluating predictions using independent observations. Predicted destructive biomass was close to measured destructive biomass for small and intermediate-sized plants, with most observations near the 1:1 line (Fig. 3A). Prediction accuracy decreased for the largest high-water plants in the later harvests, with digital biomass both over- and underestimating destructive biomass. Overlapping leaves and increased canopy complexity may contribute to reduced performance at larger plant sizes, although the cause cannot be resolved from these data. The strong relationship between digital biomass and destructive biomass observed in tobacco may also differ in species with more complex canopy architectures, such as highly branched soybean or grasses with overlapping leaves. Digital biomass measurements should therefore be validated for the species, canopy architectures, and plant sizes expected in subsequent experiments.

Prediction of destructive WUE from digital WUE was weaker than the prediction of destructive biomass from digital biomass. WUE was calculated as aboveground biomass divided by cumulative gravimetric water use, and the same water-use measurement was used in both digital and destructive WUE calculations. Therefore, differences between digital and destructive WUE arise primarily from the biomass numerator. This suggests that a biologically and statistically direct approach is to first calibrate digital biomass against destructive biomass, use the resulting model to predict biomass in an independent dataset, and then calculate predicted WUE using the common gravimetric water-use denominator.

### 4.2 Drought effects on growth and water use

Drought plants were significantly smaller than high-water plants based on both digital and destructive biomass measurements (Fig. S3) and consumed significantly less water (Fig. 5A). Despite these large reductions in growth and water use, drought did not increase whole-plant WUE or alter water use per unit destructive leaf area (Fig. S4). By the final harvest, high-water plants were more than twice the size of drought plants, demonstrating a strong effect of moderate water limitation on tobacco growth. A close relationship between biomass gain and water consumption has also been observed in *Arabidopsis thaliana* subjected to drought stress (Bhaskara et al., 2022), consistent with the approximately proportional reductions in biomass accumulation and water use observed here.

The Phenospex FieldScales allowed us to resolve temporal patterns in plant-associated water loss. Despite lower total water use in drought plants, there was no significant difference between treatments in evapotranspiration normalized by destructive leaf area (Fig. S4). Similar size-dependent patterns have been observed in Arabidopsis, where smaller plants transpired less than larger plants but differed when water loss was normalized to rosette area (de Ollas et al., 2023). In poplar, leaf area and growth rate were also strongly related to transpiration rate (Bogeat-Triboulot et al., 2019). We also observed nighttime water loss rates approximately 10% of daytime rates. Nighttime transpiration of a similar magnitude has been reported in grape (*Vitis vinifera*) (Escalona et al., 2013; Fuentes et al., 2014), whereas nighttime rates in sunflower can approach 20% of daytime rates (Rawson & Begg, 1977). Such nighttime water loss is important for whole-plant WUE because it contributes to cumulative water use during periods without concurrent photosynthetic carbon gain (Medrano et al., 2015). Continuous gravimetric phenotyping can capture these diel patterns and thereby improve estimates of integrated whole-plant water use and WUE.

### 4.3 Drought and WUE

We did not observe an increase in whole-plant WUE in tobacco subjected to moderate drought-stress. Previous work in tobacco similarly reported reductions in shoot biomass and total water consumption under short-term drought, although WUE decreased in that study (Ryan et al., 2013). Drought sensitive barley genotypes have shown little change in WUE under drought, whereas drought-tolerant genotypes increased WUE (Mansour et al., 2017), illustrating that the response of whole-plant WUE to water limitation can vary among genotypes and experiments. WUE has also been associated with plant size in Arabidopsis (de Ollas et al., 2023), whereas we did not observe a relationship between size and WUE in tobacco. Because whole-plant WUE is the ratio of biomass accumulation to cumulative water use, WUE changes only when growth and water use respond disproportionately. In this study, drought reduced both biomass accumulation and total water use without significantly changing WUE or water use per unit destructive leaf area, indicating that growth and water use declined approximately in proportion under moderate drought.

### 4.4 High-throughput phenotyping for improving estimates of plant WUE

Improving WUE has long been a target for crop breeding (Leakey et al., 2019). Intrinsic and instantaneous WUE can be estimated rapidly and nondestructively using leaf gas exchange, whereas whole-plant WUE requires integration of biomass accumulation and water use over longer periods. Leaf-level measurements are often made on fully expanded, well-lit leaves during portions of the day when photosynthesis, stomatal conductance, and transpiration are relatively high (Medrano et al., 2015) and therefore do not capture leaf and plant respiration or changes in transpiration throughout a twenty-four hour period, particularly at night. Phenotyping platforms that combine repeated imaging with gravimetric measurements can reduce this temporal limitation by providing repeated estimates of biomass and continuous measurements of water use, including day-night variation. These systems also reduce manual effort and can support greater replication or screening of large diversity panels. However, phenotyping platforms are expensive, technically complex, and their digital outputs require validation across the species, canopy architectures, and plant sizes for which they will be used. Our results demonstrate strong performance of digital biomass estimates across much of the measured range but reduced accuracy in the largest plants underscoring the importance of defining the operating range over which digital traits are reliable.

## 5 Conclusions

Digital biomass provided a strong proxy for destructive biomass across much of the measured plant-size range, with similar relative variability between methods and good predictive performance, although accuracy declined for the largest plants. Digital and destructive WUE also had similar coefficients of variation, but the weak direct relationship between the two WUE estimates indicates that additional calibration and validation are needed. Moderate drought reduced tobacco biomass and total water use without significantly changing whole-plant WUE, indicating that growth and water use declined approximately in proportion. Combining repeated digital estimates of biomass with continuous gravimetric measurements of water use provides a promising framework for resolving whole-plant WUE through development.

## AUTHOR CONTRIBUTIONS

**Samantha S. Stutz:** Conceptualization; performed the experiment and analyzed data; writing— original draft; writing—review and editing. **Ron Edquilang:** Maintained equipment; performed the experiment and analyzed data, writing—review and editing. **Carl J. Bernacchi:** Conceptualization; writing—original draft; writing—review and editing. **Donald R. Ort:** Conceptualization; writing—original draft; writing—review and editing.

## ACKNOWLEDGEMENTS

We thank J. McGrath for helpful discussion and statistical advice. We thank M. Altschuler for critical review of the manuscript. This work was done under the Realized Increased Photosynthetic Efficiency (RIPE) project supported by Gates Agricultural Innovation grant investment ID 57248.

## SUPPLEMENTAL MATERIAL

**Fig. S1.** The Phenospex FieldScan setup at the University of Illinois High Throughput Phenotyping Facility in Champaign, IL, USA.

**Fig. S2.** Layout of the drought and high-water plants for the experiment.

**Fig. S3.** Wet destructive and digital biomass across all the harvests.

**Fig. S4.** Destructive and digital leaf area across all the harvests.

**Fig. S5.** The ratio of evapotranspiration to destructive and digital leaf area.

