## Supplemental figures for "Destructive harvest validation of high-throughput measurements show that water use efficiency is unaffected by moderate drought in tobacco"

**Supporting Figure 1**

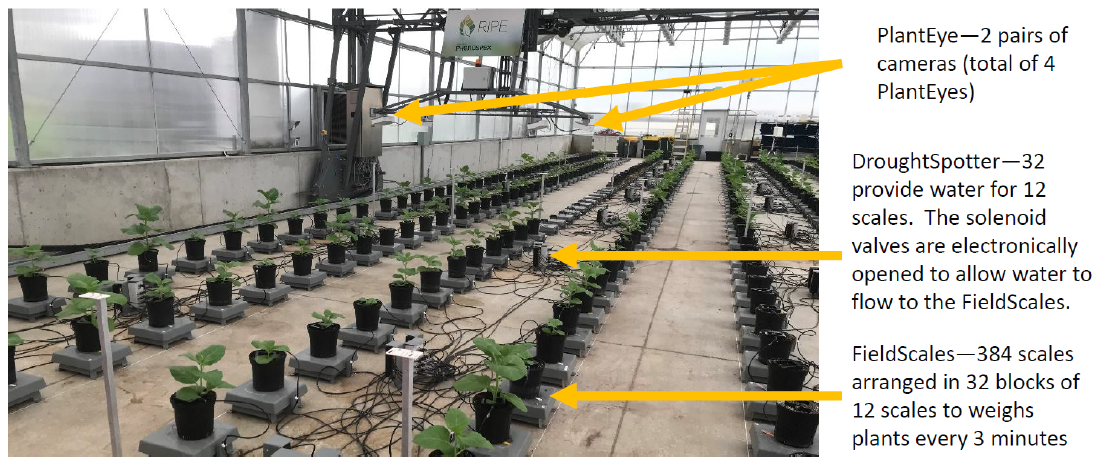

**Supporting Fig 1.** The Phenospex FieldScan^®^ setup at the University of Illinois High Throughput Phenotyping Facility in Champaign, IL, USA. The Phenospex, is composed of the PlantEye, FieldScales, and DroughtSpotter.

**Supporting Figure 2**

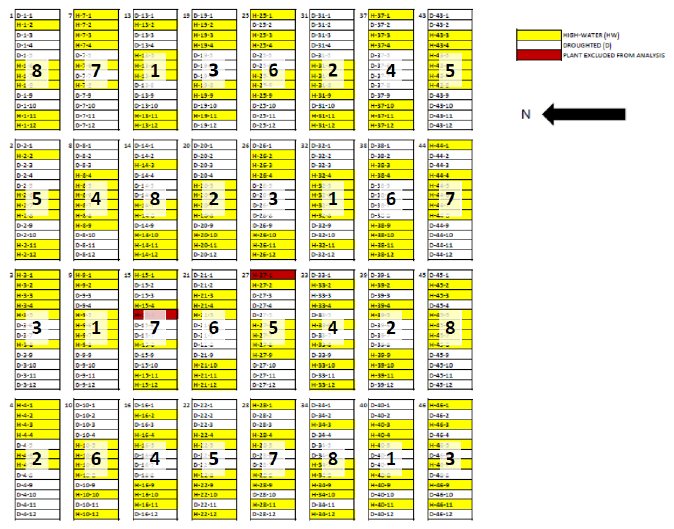

**Figure S2.** Layout of the drought and high-water plants for the experiment. Two plants were excluded from the final analysis. Numbers in the middle indicate when the blocks were harvested.

**Supporting Figure S3**

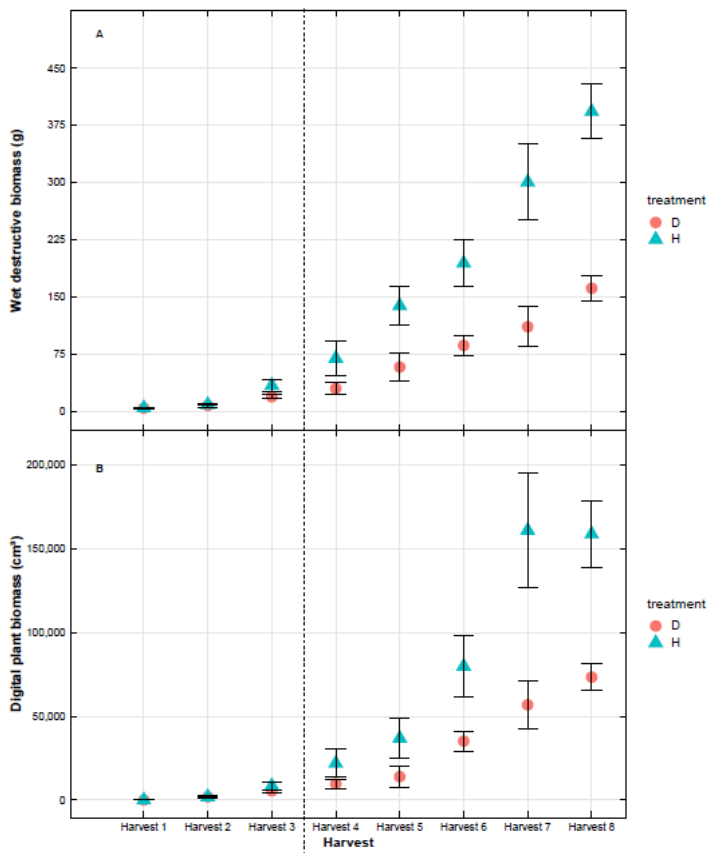

**Figure S3. a)** Wet destructive biomass across all the harvests. **b)** Digital plant volume across all the harvests for tobacco plants grown under drought conditions (D—41% soil field capacity) and high-water (H—65% soil field capacity). Error bars represent ±1 SD of four replicates for each harvest. The dotted line represents the start of the drought experiment, where more than 50% of the drought plants were watered.

**Supporting Figure S4**

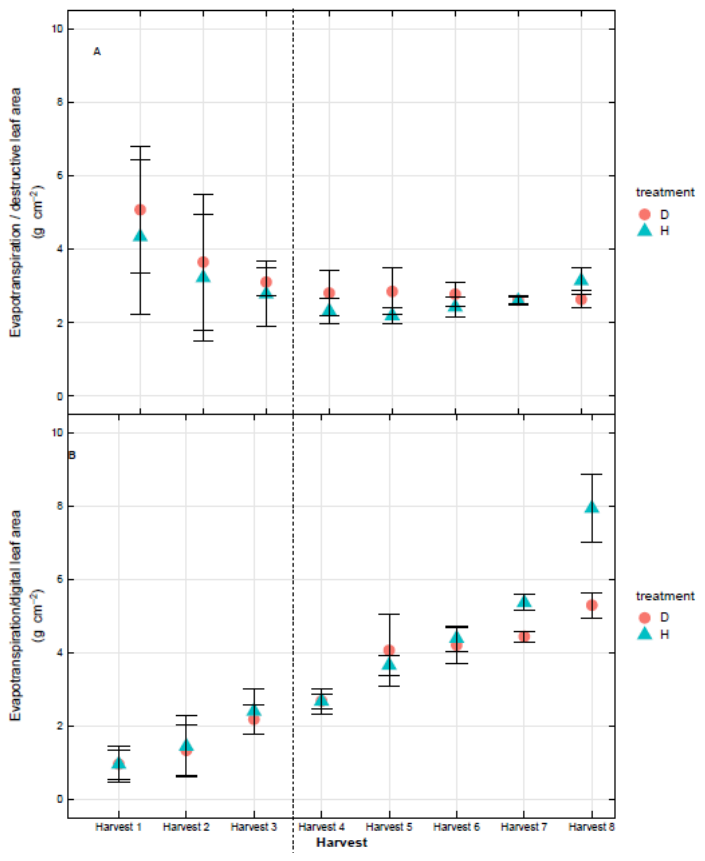

**Fig. S4.** A) The ratio of evapotranspiration destructive leaf area and B) evaporatranspiration to digital leaf area across all the harvests for tobacco plants grown under drought conditions (D—41% soil field capacity) and high-water (H—65% soil field capacity). Error bars represent ±1 SD of four replicates for each harvest. The dashed line showed the start of the drought experiment, where at least 50% of the remaining drought plants were watered.

**Supporting Figure S5**

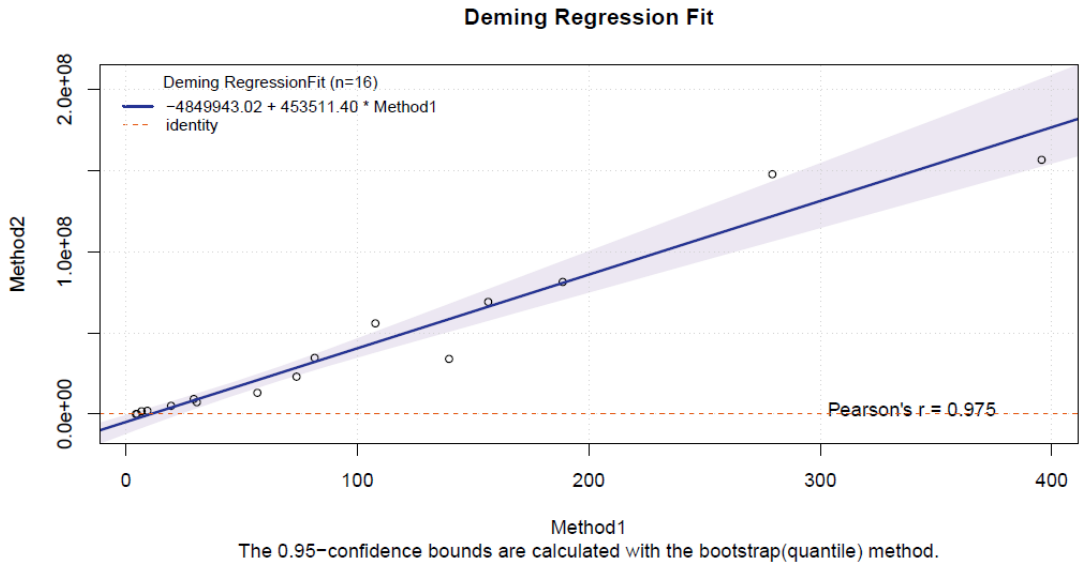

Wet destructive biomass (g)

Digital biomass

Wet destructive biomass (g)

**Figure S5.** Deming Regression model fit to half of the biomass data. The model indicates there is a high correlation between digital and destructive biomass.

**Supporting Figure S6**

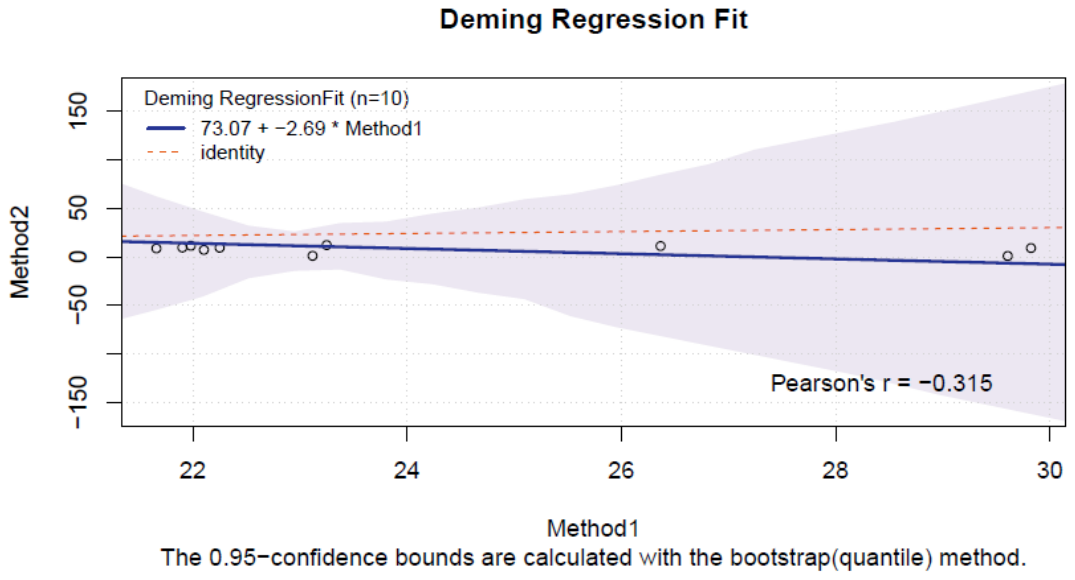

Digital WUE

Destructive WUE

Destructive WUE

**Figure S6.** Deming Regression model fit to half of the WUE data. The model indicates there is a moderate correlation between digital and destructive WUE.

**Supporting Figure S7**

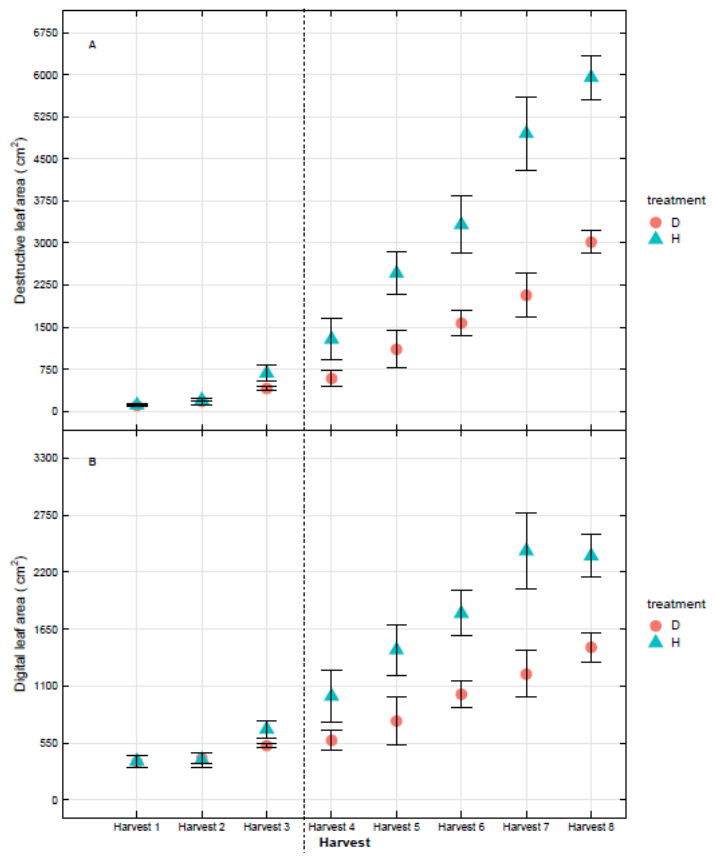

**Fig. S7.** A) Destructive leaf area and B) digital leaf area across all the harvests for tobacco plants grown under drought conditions (D—41% soil field capacity) and high-water (H—65% soil field capacity). Error bars represent ± 1 SD of four replicates for each harvests. The dotted line represents the start of the drought experiment, the point where over 50% of the remaining drought treatment plants were watered.
